# Defining Network Topologies that Can Achieve Molecular Memory

**DOI:** 10.1101/2024.03.13.584853

**Authors:** F. Sevlever

## Abstract

In the context of cellular signaling and gene regulatory networks, the concept of molecular memory emerges as a crucial determinant of molecular mechanisms. This study introduces a novel memory quantifier designed to comprehensively capture and quantify the memory of a system in response to transient stimuli. We proposed and validate this quantifier through toy models, showcasing its effectiveness in systems with positive feedback loops and bistability. In addition, we develop an algorithm to assess long-term memory in circuits, leading to the identification of minimal motifs that play pivotal roles in conferring memory.

The research explores the comparative impact of positive and negative feedback loops on memory, revealing that positive feedback enhances memory while certain negative feedbacks may diminish it. An intriguing finding emerges as oscillating circuits, even in the absence of positive feedback, exhibit memory, with the phase of oscillations storing information about stimulus duration.

Finally, we experimentally validate the quantifier using mouse Embryonic Stem Cells (mESCs) subjected to transient differentiation stimuli. The proposed memory quantifier is applied to gene expression dynamics, revealing varying degrees of memory retention among different genes. The vectorial nature of the quantifier proves advantageous in capturing the holistic memory dynamics of the system.

## Introduction

In the context of cellular signaling and gene regulatory networks, there is an increasing evidence that memory can be a key determinant factor underlying molecular mechanisms (1)(2)(3)(4) (5)(6)(7)(8). In these papers, cellular systems are first treated with *priming* inputs followed by a time window and then the stimuli. The observation is that response to the stimuli is dependent of the priming inputs, so each system can remember it for at least the duration of the time window.

From a general perspective, memory is usually defined in a context dependent manner. In computing, memory is the amount of information that some device can store. In complex systems, the final state or position dependency with its history; the bi or multi-stability or the system hysteresis. In molecular biology, we found two main definitions. One is the capacity of past stimuli information storage (9)(10); the other is the stationary response to a transient stimulus (11).

A priori, these definitions can appear different, but they have common fundamentals. A computer memory device, like a hard drive disk of 1TB, can store information in magnetic cells. These cells are magnetized with an external magnetic field, and they are capable of keep a stationary magnetization even when the transient external field have passed long ago (12). The magnetization value is dictated by the history of external field. Dynamically, it has two stable fixed points, and the external field moves the magnetization from one to another. Hysteresis of this system arise from bistability, which is the property of a system having two stable fixed points.

It is well known that positive feedback loops can provide bistability, hysteresis and memory (3)(5)(6)(10)(13)(14)(15)(16)(17)(9)(18). Imagine a molecule that, above certain concentration, turn on its own synthesis. If there is not sufficient concentration, our molecule will decay to zero, reaching a fixed point with low or no concentration at all. But, if we could give a transient stimulus that activates its synthesis until the threshold concentration is exceeded, then the molecule would remain turning on itself into another stable fixed point, with high concentration.

The previous different definitions of memory, even when the fundamental concept is similar, result in different ways of measuring it, which then cannot be comparable. For example, in molecular biology, studies are often oriented to prove if some system has or not memory, in a binary way, and is impossible then to determine if some signaling pathway remember one stimulus more than another. Even further, how can we compare memory of different pathways where they have not the same number of molecular actors?

In this work, we aim to give a general mathematic and unique definition of the memory of a system, and the associated measurable observables to quantify it. We first propose this general definition for dynamical complex networks as our most general approach. Then, we computationally test it for all the possible 1, 2 & 3-nodes circuits, asking which are the motifs that have memory and what are their underlying mechanisms. Finally, we measure memory in mouse embryonic stem cells Gene Regulatory Network and discuss some other examples.

## Results

### Proposed memory quantifier

We propose a quantifier of the memory a system has from a received stimulus 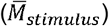, with 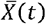 representing the state or of the system at a given time *t*:

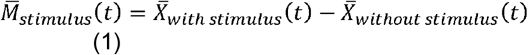

In Fig. 1 we illustrate how to compute 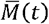 for a toy model. This model has two nodes A and B, both have basal activation and inhibit one another in a symmetric way (Figure 1A up). This double inhibition works as positive feedback (if A inhibits B who inhibits A back, A activates itself). In general, positive feedback can provide bistability, and there are plenty of examples in molecular biology (19). Here, one stable fixed point (green circle in Figure 1A bottom) corresponds to A high inhibiting B, which remains low, and the opposite where B is high and A low. The phase diagram shows the trajectories departing from different initial conditions in the border of the phase space and reaching one fixed point or another.

**Figure 1:**
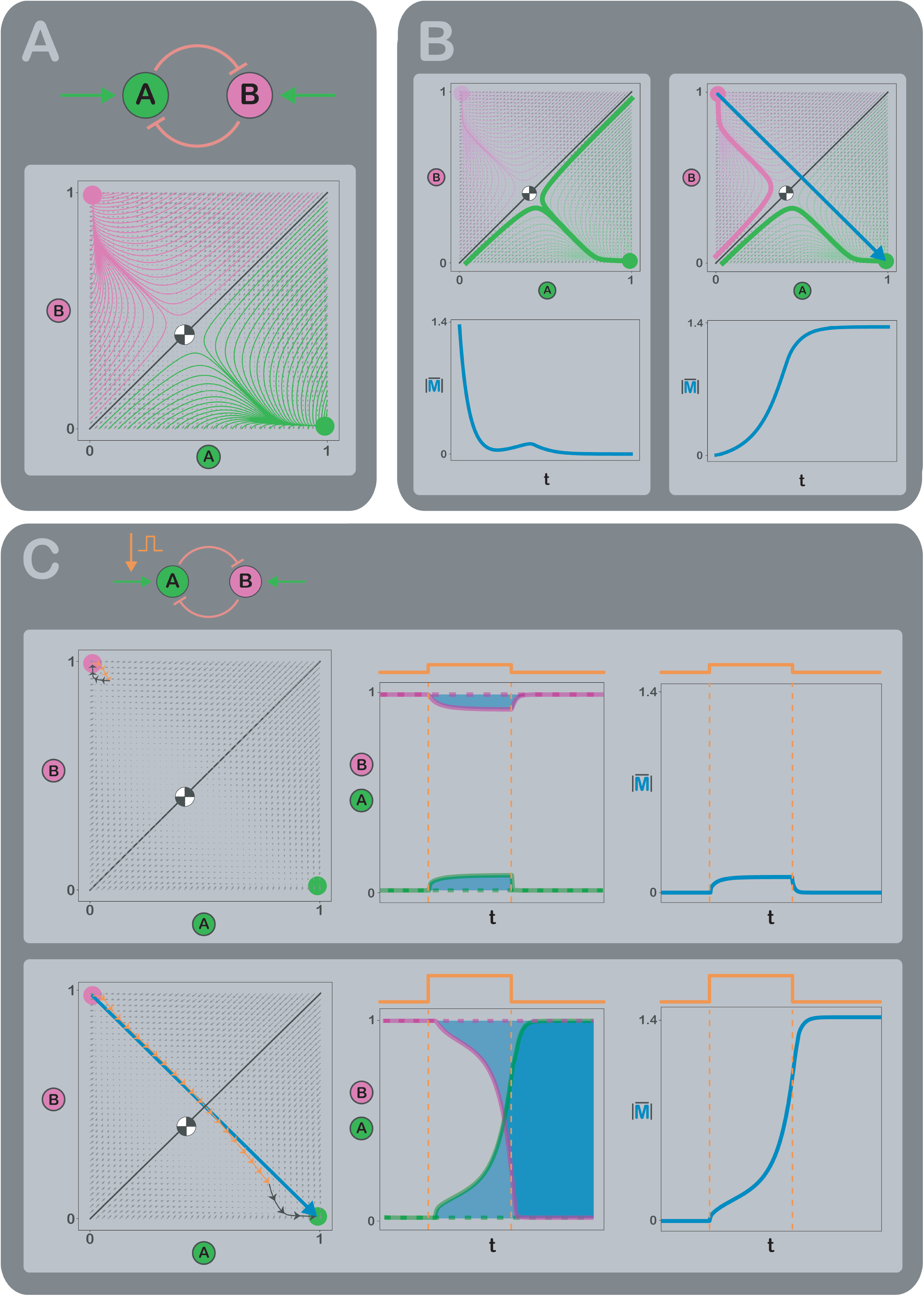
Circuit multistability and memory. A. Example circuit with two nodes inhibiting each other symmetrically (top). This positive feedback loop provides bistability. The phase diagram (bottom) shows several trajectories in magenta and green departing from the edges and reaching the two fixed points. The magenta circle is the stable fixed point where B node “wins” and A node “loses”. The green circle is the opposite. As both inhibitions are symmetrical, the node of higher initial concentration determine the final fixed point of the trajectory. The diagonal black line indicates this separation and the black and white circle is the fixed unstable point. Background grey arrows are the velocity field. B. Two examples of persisting information from initial state. Top left: two trajectories depart from opposite states and reach the same fixed point. Bottom left: memory module, as the distance between both trajectories in time, goes to zero. Measuring the final state does not provide information of the initial state. Top right: two trajectories depart from almost same state and reach different fixed points. Bottom right: memory module remains positive even when *tH ∞*. Measuring the final state does provide information of the initial state. C. Same dynamic examples of panel B but replacing initial conditions for square pulse stimuli. Low (top) and high (bottom) pulse stimulus produce different permanent responses. Phase diagram trajectories (left column) are divided in three stages: before (black arrowed line), during (orange arrowed line), and after (black arrowed line) stimulus. On each stage, system evolve until steady state is reached. Low stimulus is not sufficient to move the state through the black line from the magenta fixed point to the green one. On contrary, high stimulus is enough and the circuit change state permanently. Temporal curves for A and B nodes concentrations are in the middle column. Continuous lines for stimulated curves and dashed lines for non-stimulated curves. Notice the interchange between both nodes concentrations for high stimulus. Right column shows memory module in time. While low stimulus goes to zero, high stimulus remains in a positive value.

Figure 1B computes the temporal evolution of the distance between two pairs of trajectories. Notice that these two pairs are opposite examples of past information storage. In one case, the system starts at two very distant points in the phase space and ends in the same fixed point. It is not possible to identify which was the initial condition once the system has reached the fixed point. The two trajectories collapse into the same one. In the opposite case, both trajectories start at very similar initial conditions but their distance increases in time and then remain high when they end up in different fixed points. If we measure A or B value, we could discriminate which was the initial condition.

This example points out that bistability provides memory to a system, in the way that each fixed stable point stores some information about the past. The blue arrow is 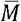 at the final time. In this example we think of two different initial conditions. But, in general, any transient stimulus can be thought of as a variation in the initial condition of a system. In this line, Figure S1 shows some interpretations and mathematical properties of out quatifier.

Figure 1C illustrates how to measure 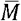 in a practical way, using a transient square pulse stimulus given to the basal activation of A (Figure 1A up). Left panels compare two trajectories in the phase space for low and a high amplitude stimulation. The high amplitude stimulus is sufficient to move the system from one fixed point to the other while the low one is not. Middle panels show the time courses of A and B and the distance between them 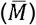. Right panels show 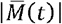 and how it is different from zero only if the stimulus is sufficiently high so that the system can remember it.

In conclusion, we propose 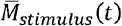 as a quantifier of the memory a system has from a transient stimulation. It is a vector with the dimension of the system. In our example, it has dimension 2. Each coordinate of 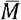 is the difference of the value of each variable with and without the stimulus. If 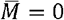, the state of the system is equal with or without the stimulus, so it corresponds to no memory. Notice that each 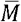 coordinate can be positive or negative. This accounts that the system can remember a stimulus no matter if it activates or inhibits some variable.

### Algorithm to determine if any circuit has long-term memory

The aim of this section is to design an algorithm to measure the memory of any circuit and then use it to find mechanisms that can provide memory to a system.

We first tested 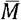 on another two toy models with only one node. One of them has a well characterized memory while the other does not. Both have basal activation and inhibition, and the difference is that one of them has a positive autoregulation (Figure 2B, left). Autoregulation provides positive feedback leading to bistability and memory, as shown in the previous section. The bistability depends on the relative strength of the feedback and the basal inhibition, which is given by the parameters.

**Figure 2:**
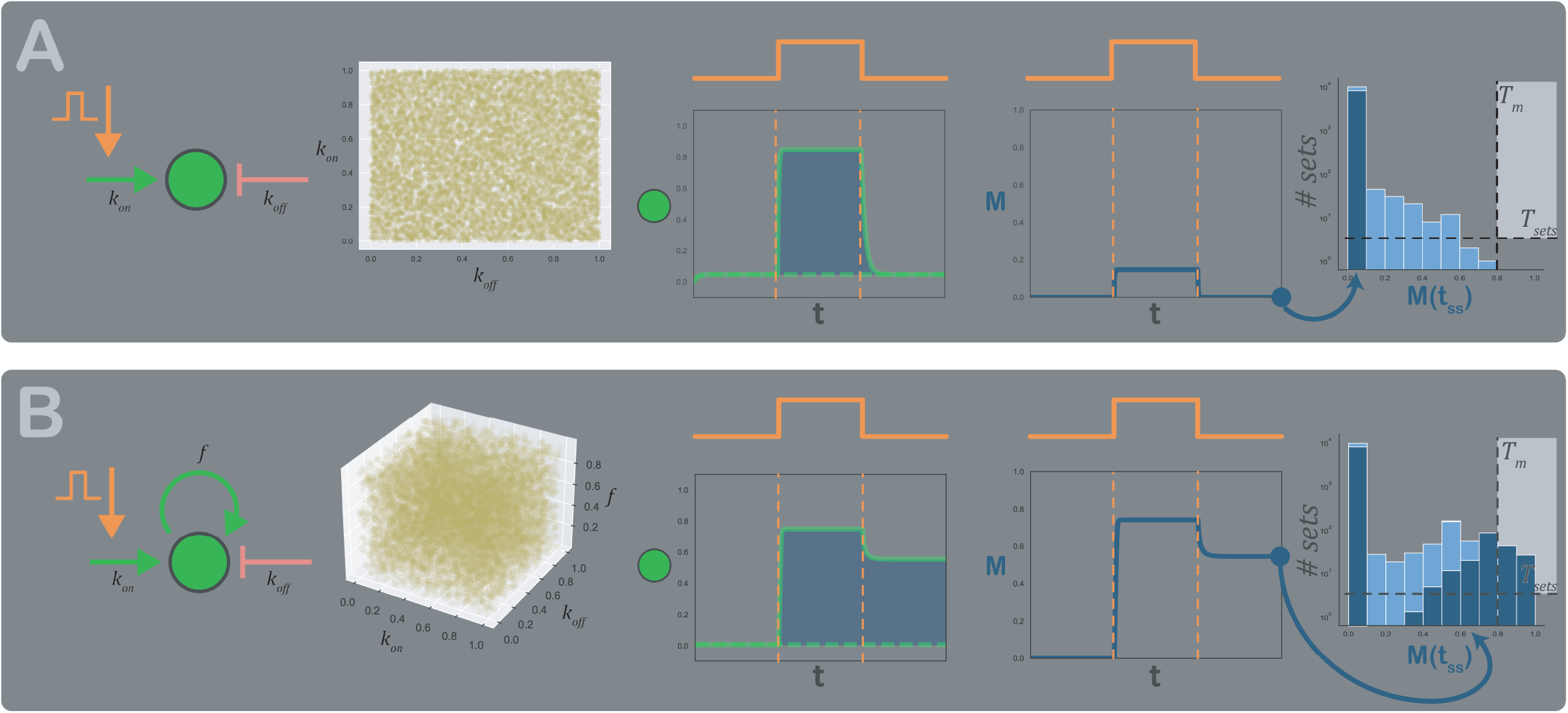
Algorithm to determine if any circuit has long-term memory. Algorithm application to two example topologies (left). Panel A topology is the simplest with one node and has only one stable fixed point. Panel B topology has an autoregulation and could have bistability. The algorithm consist in take 10000 random sets of parameters (center-left) and integrate the circuit for each one with and without the stimulus (center). Then compute memory vector in time (center-right) using eq. (5). Finally, take the long-term memory (dark blue circles) for each set and make the histogram distribution for every set (right). Dark blue bars correspond to completely integrated circuits and light blue to interrupted ones. Light blue bars are on top of dark blue bars, not behind. Dashed lines represent both *T*_*M*_ and *T*_*sets*_ and light grey the region above these thresholds. Circuit in panel B contain bars in this region, above *T*_*M*_ whose values are greater than *T*_*sets*_. Therefore, this topology has long-term memory. On contrary, panel A topology lacks bars in the light grey region, even considering light blue bars, and then it has no long-term memory.

We computationally integrated these models with and without a square stimulus, for 10000 sets of parameters for each one using Latin Hypercube Sampling (20) (Figure 2A-B, center-left). Then, we computed 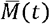 (Figure 2A-B, center) and saved, for each model and for each set of parameters, 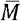 at final time (Figure 2A-B, center-right). Finally, we analyzed the distribution of 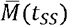 (Figure 2A-B, right). We analized the final value of 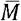 because we are interested in classifying circuits according to their long-term memory, understood as the situation in which the system never returns to its previous stimulus state.

As expected, we obtained that the model has no memory for all the analyzed parameter sets. In contrast, the model with autoregulation has some parameter sets with memory. These sets are those with the feedback sufficiently strong to give bistability but also sufficiently weak to remain “off” after the stimulus has passed.

Histograms in Figure 2A-B right, show the two thresholds we choose to define if a circuit has memory or not. A circuit having memory means that has more than *T*_*sets*_ where each one has more than *T*_*M*_.

### 1, 2 & 3-nodes mechanisms that provide memory

In this section we aim to characterize every 1, 2 & 3-nodes circuits in their ability to exhibit memory. The main goal is to find all possible mechanisms or motifs leading to long term memory. We followed the approach presented in Ma et al (21) and used in several works (22)(23)(24)(25)(26)(27)(28). This strategy considers all possible circuits (with 1, 2 and 3 nodes), with each node activating, inhibiting, or not interacting with the others and with itself (see Methods). In those cases where a node has no activation or inhibition, a basal one is provided. The total numbers of circuits are 3, 54 and 16038 for 1, 2 and 3 nodes, respectively.

Figure 3A illustrates the three sets of circuits. For each circuit, we applied the algorithm explained in the previous section (Figure 3B, left) and compute 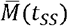 (see Methods). The pulse stimulation is applied to the green node. Figure 3B, right, shows the data obtained for a particular 3-nodes circuit. Each row contains the values of 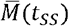, a three-dimensional vector, for a given parameter set. Many parameter sets do not retain the stimulus, but in cases where such sets exist, a thorough exploration of 10,000 has proven to be sufficient for discovering them.

**Figure 3:**
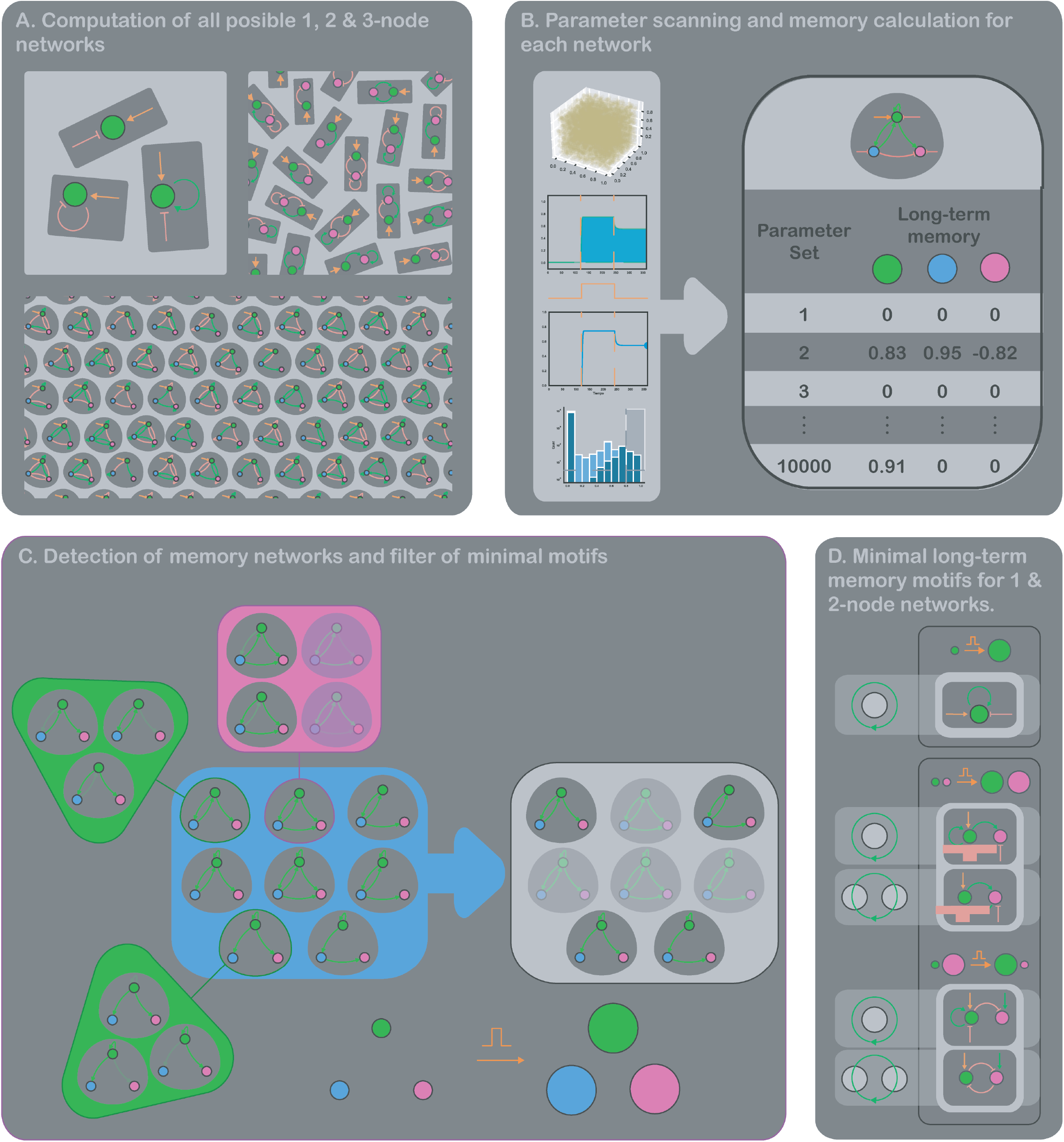
Long-term memory motifs detection. A. Considered circuit topologies representation for 1, 2 and 3-nodes circuits. The 3 possibilities for 1 node are showed, but only a small subset of all 2 and 3-nodes circuits (54 and 16038 in total). B. Long-term memory computing. For each topology, we applied the algorithm of Fig. 2 and constructed the database of memory vector for the 10000 sets of circuits. C. Motifs detection and filtering. Blue square represents a subset of the set of topologies with long-term memory in the three nodes (the three values of memory vector where above thresholds). The aim of the filter is to suppress the redundancy in motifs. Take the remarked topology in pink at top-center. The pink square has every topology combination of subtracting only one interaction. The transparent topologies are part of the original blue subset and then we discard the pink topology. Notice the same procedure applied to both green remarked topologies result in no repeated topologies. The filter takes only the not redundant topologies, which are showed in the grey square for the blue subset. D. Minimal long-term memory motifs for 1 and 2-node circuits. 1-node minimal motif is just one of the three possibilities (top square). Memory is due to the positive autoregulation. For 2-nodes (bottom square), there are two motifs where both nodes activate to record the stimulus (top pair of topologies) and two where one activates and the other inhibits (bottom pair of topologies).

After integrating each circuit for 10000 parameter sets, an algorithm for minimal motifs identification was applied (Figure 3C and Methods). The aim of this algorithm is to find the motifs responsible for a given property (in this case, long-term memory), given a group of circuits having that property. For example, suppose we want to identify circuits in which the stimulus activates all the nodes, and they remain activated. This condition means that the three coordinates of 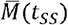 are positive. Taking the data of Figure 3B, we can fix two thresholds and filter all circuits where more than *T*_*sets*_ sets of parameters have each coordinate of 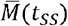 above *T*_*M*_ (see Methods). An example of the circuits obtained is showed in Figure 3C. Notice that many circuits are similar, and they all have one or more positive feedbacks, but some of them are combinations of simpler circuits. The aim of this algorithm is to avoid the redundancy of founding the same responsible motifs in different circuits, identifying only the simplest fundamental mechanisms, as shown in Figure 3C.

All 1 & 2-nodes minimal motifs with long term memory are illustrated in Figure 3D. There is only one minimal motif with 1-node having memory. As expected, it is the one with a positive autoregulation. For 2-nodes circuits, we found 4 minimal motifs having memory, classified in 2 groups: one where the stimulus activates both nodes and the other where it activates one and inactivates the other. A given node can store memory of a stimulus by an irreversibly activation (OFF to ON) or inhibition (ON to OFF). The resulting 4 minimal motifs are all the possible positive feedback involving the stimulated green node. The mechanisms 1 and 4 have been founded in developmental gene regulatory circuits. The first corresponds to a stimulus starting an irreversible development program (3) and the second to a cell making a binary decision, as it works like a toggle switch (8) (29) (30). Notice there is a perfect analogy between the two groups, where we can make a bijection from one to another. We just take a motif and change the kind of arrow (activation or inhibition) involving the regulations to or from the node that switches from activation to inactivation to record stimulus. This suggest that topologies that store memory come in opposite pairs: one where the stimulus irreversibly activates a node and another one where irreversibly inactivates it.

The results for the minimal motifs in 3-nodes circuits are summarized in Figure 4. All of them have positive feedback loops involving the stimulated node. We found 36 motifs and classified them in 4 groups according to the activation and inhibition of the same nodes, as before. The relative location of each motif into its own panel indicates similarities between motifs from different panels. Notice that motifs sharing relative location have the same regulations, leaving aside its sign. This supports the idea of symmetrical opposite motifs, considering that with 3 nodes we have 2 nodes to swap from activated to inactivated (blue and magenta). Then, for a given motif, we can swap just the blue node, the magenta node or both, giving four similar mechanisms. We did not find any motif where the green node inactivates. This is probably because it always receives an activating stimulus. The fact that our method found every 4 motifs of each mechanism without using this symmetry indicates robustness.

**Figure 4:**
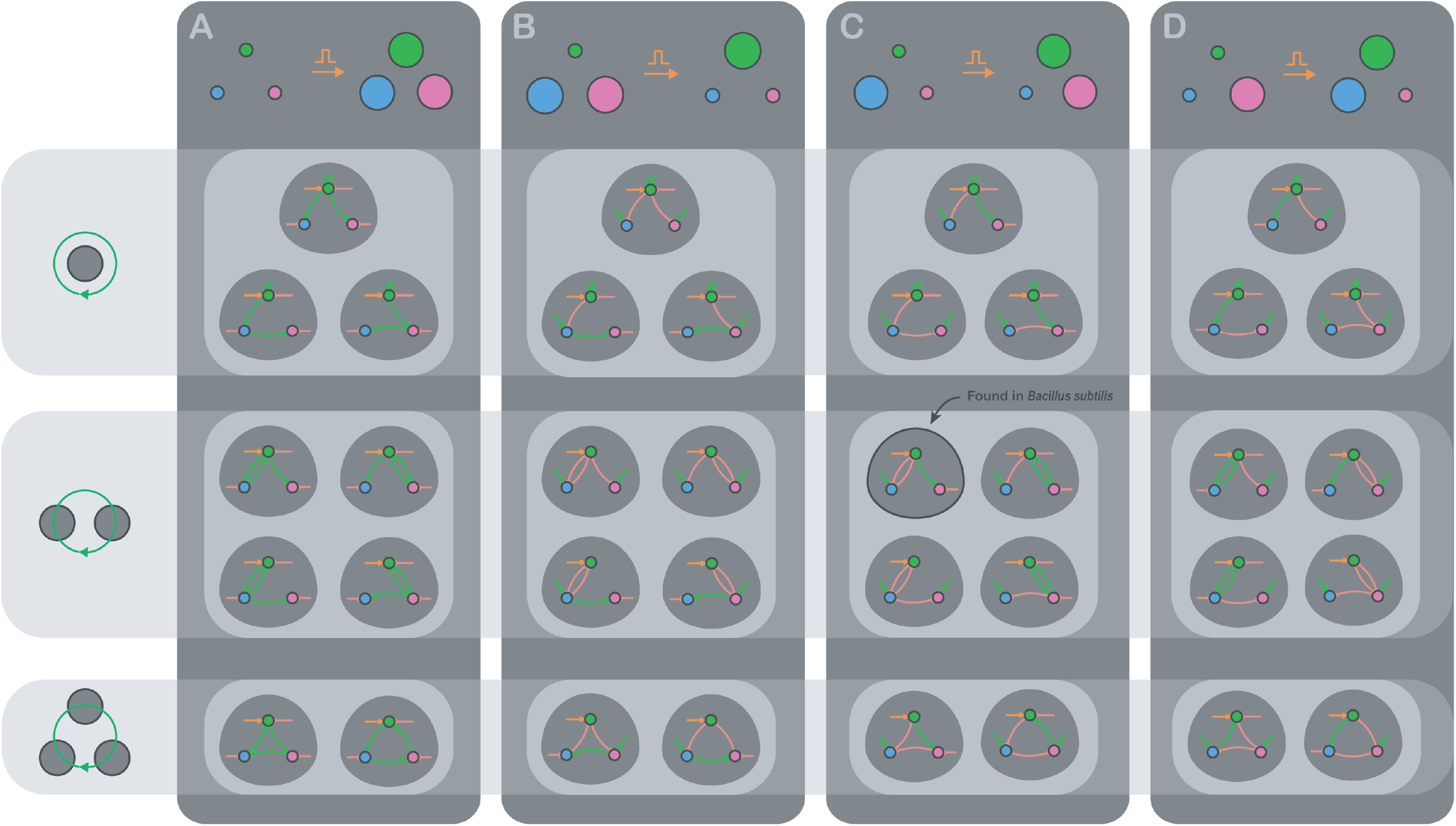
3-nodes long-term memory minimal motifs. Panel titles indicates if memory is recorded in activated o inhibited nodes. There are 48 total minimal motifs. Every motif founded is a positive feedback. Topologies are categorized in three rows for 1, 2 or 3-nodes involved in the positive feedback. Every positive feedback involve the green node, which receives the input. A. Three nodes activated. 1-node row: three similar mechanisms where the green node activates itself and the others directly (top) or indirectly (bottom). Both founded topologies at bottom are symmetrical in swapping blue and magenta nodes. 2-nodes row: double positive regulations where the remaining node is activated by the green node directly (top) or indirectly (bottom). Each topology at left is symmetrical to its right in swapping blue and magenta nodes. 3-nodes row: triple positive regulations define a rotating direction and we found both symmetrical clockwise and anti-clockwise topologies. B. Green activated, blue, magenta inhibited. 1-node row: three similar mechanisms where the green node activates itself and inhibits the others directly (top) or indirectly (bottom). Both founded topologies at bottom are symmetrical in swapping blue and magenta nodes. 2-nodes row: double negative regulations where the remaining node is inhibited by the green node directly (top) or indirectly (bottom). Each topology at left is symmetrical to its right in swapping blue and magenta nodes. 3-nodes row: double negative positive regulations define a rotating direction and we found both symmetrical clockwise and anti-clockwise topologies where the green node inhibits the others. C. Green, magenta activated, blue inhibited. Every founded topology has an analogous partner in panel D, because of the symmetry of swapping blue and magenta nodes. 1-node row: three similar mechanisms where the green node activates itself and the magenta and inhibits the blue both directly (top) or indirectly (bottom). 2-nodes row: double negative regulations (left) where the magenta node is activated by the green node directly (left-top) or indirectly (left-bottom). Double positive regulations (right) where the blue node is inhibited by the green node directly (top-right) or indirectly (bottom-right). 3-nodes row: double negative positive regulations define a rotating direction and we found both symmetrical clockwise and anti-clockwise topologies where the green node activates the magenta and inhibits the blue, directly or indirectly depending on clockwise or anti-clockwise direction. D. Green, blue activated, magenta inhibited. Every founded topology has an analogous partner in panel C, because of the symmetry of swapping blue and magenta nodes. 1-node row: three similar mechanisms where the green node activates itself and the blue and inhibits the magenta both directly (top) or indirectly (bottom). 2-nodes row: double positive regulations (left) where the magenta node is inhibited by the green node directly (left-top) or indirectly (left-bottom). Double negative regulations (right) where the blue node is activated by the green node directly (top-right) or indirectly (bottom-right). 3-nodes row: double negative positive regulations define a rotating direction and we found both symmetrical clockwise and anti-clockwise topologies where the green node activates the blue and inhibits the magenta, directly or indirectly depending on clockwise or anti-clockwise direction.

Between panels B and C, there is another bijection between motifs due to the symmetry of swapping the blue and magenta nodes. This symmetry does not exist in case of 1 and 2-nodes circuits because the green node receives the stimulus and therefore is not equivalent to the other nodes. Our method found again the motifs without using this symmetry, another proof of its robustness.

The minimal motif remarked in Figure 4C has been found in *Bacillus subtilis*, responsible for differentiation to a transient competent state (31).

Summarizing, every minimal motif found is a positive feedback. In this line, the next question we investigate is if every kind of positive feedback improves the memory of a given circuit.

### Neighboring circuits comparative algorithm: positive feedbacks improve memory, and negative feedbacks erase it

In this section, we aim to compare every 2 & 3-nodes circuits to answer if those with positive feedbacks have more memory than those without. Although, we want to quantify this comparison to measure if different feedbacks are better providing more memory than others.

We constructed the space of circuits. A meta-network for each set of 1, 2 & 3-nodes circuits containing all of them. This is a directed network with two kinds of edges: activation or inhibition. The construction follows two simple rules: 2 circuits are neighbors if you can depart from one and reach the other just adding or removing one regulation arrow, and the direction of the edge is always in the addition way. If you added an activation of one node to another, the edge arrow is green, or red if the added regulation was an inactivation (Figure 5A). Because there are 9 possible regulation arrows for the 3-nodes circuits, its meta-network has 9 possible directions. The topology of this network is the same as a hypercube of 9 dimensions. Each dimension has large 3 as the arrow can be an inhibition, no regulation, or activation.

**Figure 5:**
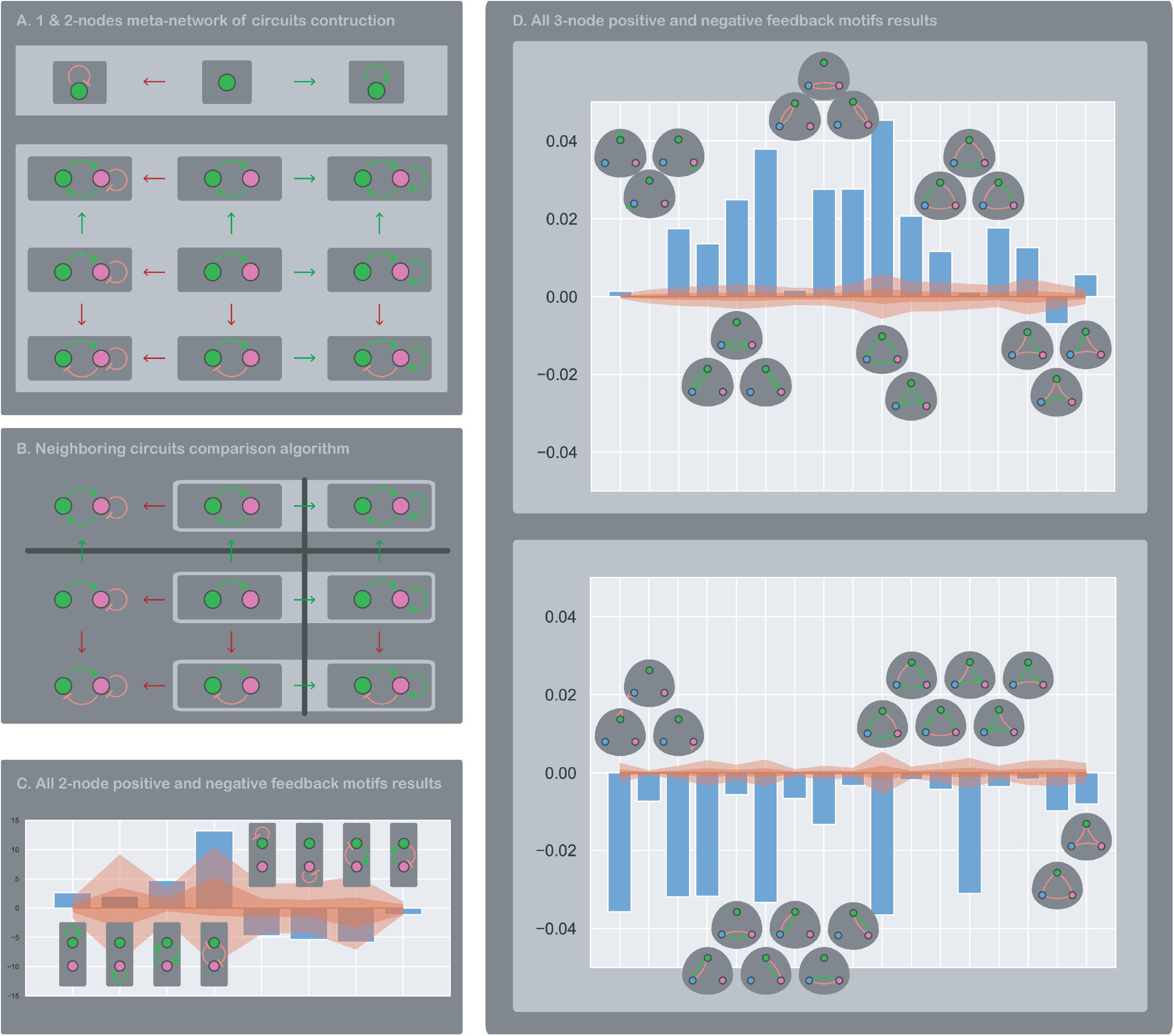
neighboring circuits algorithm. A. 1 & 2-nodes meta-network of circuits construction. The total three 1-node circuits ordered in the meta-network (upper panel). As there is only one possible edge, the autoregulation of the unique node, the meta-network has only one dimension. In this direction, the green arrow pointing to the left indicates adding an activation edge and the red arrow pointing right, an inactivation edge. The network shows the central circuit with no edges and its two directed neighbors with the autoactivación (right) and autoinhibition (left). Bottom panel shows a part of the 2-node circuits meta-network, for the neighborhood sharing the magenta node autoregulation (horizontal direction) and the regulation edge from the magenta to green node (vertical direction). Going right means to add an autoactivation to magenta node; left, to add an autoinhibition; up, to add an activation from magenta to green node and down, an inhibition. B. Neighboring circuits comparison algorithm. An example of how the algorithm determines if a motif provides memory. Here, the example motif has one edge, the autoactivation of magenta node. The first step is cutting the meta-network through the line perpendicular to the direction of the addition of one of the motifs edges (an hyperplane in the complete meta-network). In this example, this line is the vertical black line passing through green arrows pointing right, in the direction of the addition of the magenta autoactivation. Then, the pairs of neighbor circuits are the ones from one and another side of this line (resalted in light grey), which differ by just this autoactivation. The final step is to compute the difference in memory into each pair and then the mean over every pair (eq. (6)). If our motif has more than one edge, this idea can be extended by cutting with one hyperplane for any edge our motif has. In our example, if the motif now has the autoactivation plus the activation from magenta to green node, we add the horizontal black line which passes through adding this second edge direction. The only remaining circuit with the motif is the one in the upper right corner, and the resulting neighbor pairs are both crossing each line. In the complete meta-network, the lines are hyperplanes, and the pairs are the ones along each corner. C. All 2-node positive and negative feedback motifs results. Results of the application of the neighboring circuits comparative algorithm to every positive and negative feedbacks motif in the 2-nodes circuits. Each bar corresponds to eq. 6 computation of each motif indicated. Light and dark red bands show control tests 95% and 50% confidence intervals. Every positive feedback has a bigger or similar bar than its 95% control band, and every negative feedback, a lower or similar. This implies that positive feedbacks, in general, improve the memory of its circuits, and negative feedback, erases it. D. All 3-node positive and negative feedback motifs results. Results of the application of the neighboring circuits comparative algorithm to every positive and negative feedbacks motif in the 2-nodes circuits. Each bar corresponds to eq. 6 computation of each motif indicated. Light and dark red bands show control tests 95% and 50% confidence intervals. Almost all positive feedbacks have a bigger or similar bar than its 95% control band, and almost every negative feedback, a lower or similar. This implies that positive feedbacks, in general, improve the memory of its circuits, and negative feedback, erases it.

The meta-network allows us to define distances and regions of some similarities between circuits. Suppose we want to take all 2-nodes circuits where the magenta node has a positive autoregulation. This operation consists in just drawing a plane cutting the hypercubic network perpendicularly to its arrows for the addition of the desired autoregulation. Figure 5B illustrates this process, where every circuit at the right side of the vertical plane has the regulation and no one at left does. Notice that the plane divides pairs of neighboring circuits. In each pair, both circuits are equal except for the regulation chosen, and then are comparable. The Neighboring Circuits comparative algorithm is based on comparing these circuits (see Methods) to determine if a given regulation improves or not the memory of a circuit. If we want to determine this but for a given motif, composed by more than one regulation, it is possible to extend this idea. Suppose our motif is again the autoregulation of the magenta node plus an activation from this node to green node. This motif is composed by two regulations, and we need to draw a plane for each one. The region with our motif will be just the intersection of every resulting region having each regulation. In Figure 5B, it is just the circuit at top right, because it is the intersection between the right column and the top row of the network.

We applied this algorithm to every positive and negative feedback motif for the 2 & 3-nodes meta-networks. The results are illustrated in Figure 5C for 2-nodes positive and negative feedbacks and Figure 5D for positive (upper panel) and negative (bottom panel) feedbacks. The main conclusion is that positive feedbacks provide memory to a circuit while some negative feedbacks can take it out. This last observation is due to negative feedbacks shorten the dynamic range. Imagine that a molecule inhibits its own synthesis, then, for a given basal constant synthesis rate, its stationary state will be lower compared to no autoinhibition. This molecule will never explore higher values, shortening its dynamic range. The link to memory is that, if some circuit has bistability, a negative feedback could shorten the dynamic range explored after some stimulus, keeping the circuit away from the other fixed stable point. The circuit will no longer remember the stimulus. In this line, it is remarkable that negative feedbacks, which are better to remove memory, are those where the stimulated node (green) is directly inhibited. On the opposite direction, positive feedbacks that are better for memory are those where the output magenta node is directly activated.

### Oscillations can store memory of a stimulus

So far, we have seen that positive feedbacks can provide memory. And this arises always from bi or multi-stability, where the stimulus carries the circuit from one fixed stable point to another (Figure 1). In this section, we ask if multistability is the only cause of memory, or if exist another dynamical feature that could also provide memory.

We apply our memory quantifier to the simplest circuit that can oscillate (Figure 6A, upper panel). The oscillation is due to a mechanism called frustrated bistability (32). This kind of mechanism has been founded in nature (33) and presents robust oscillations arising from the coupled positive and negative feedbacks. The positive feedback provides the hysteresis which supplies the required delay between both nodes to have oscillations. This circuit has memory because a stimulus can turn self-sustained oscillations on (33). This mechanism accomplishes to remember a stimulus while it lacks two different stable fixed points. The stimulus rides the system from a stable fixed point to a limit cycle, where it remains oscillating. However, the circuit has hysteresis and some kind of bistability, given by its positive feedback. So, we redirect our original question: is there any oscillatory circuit with memory that lacks any positive feedback?

**Figure 6:**
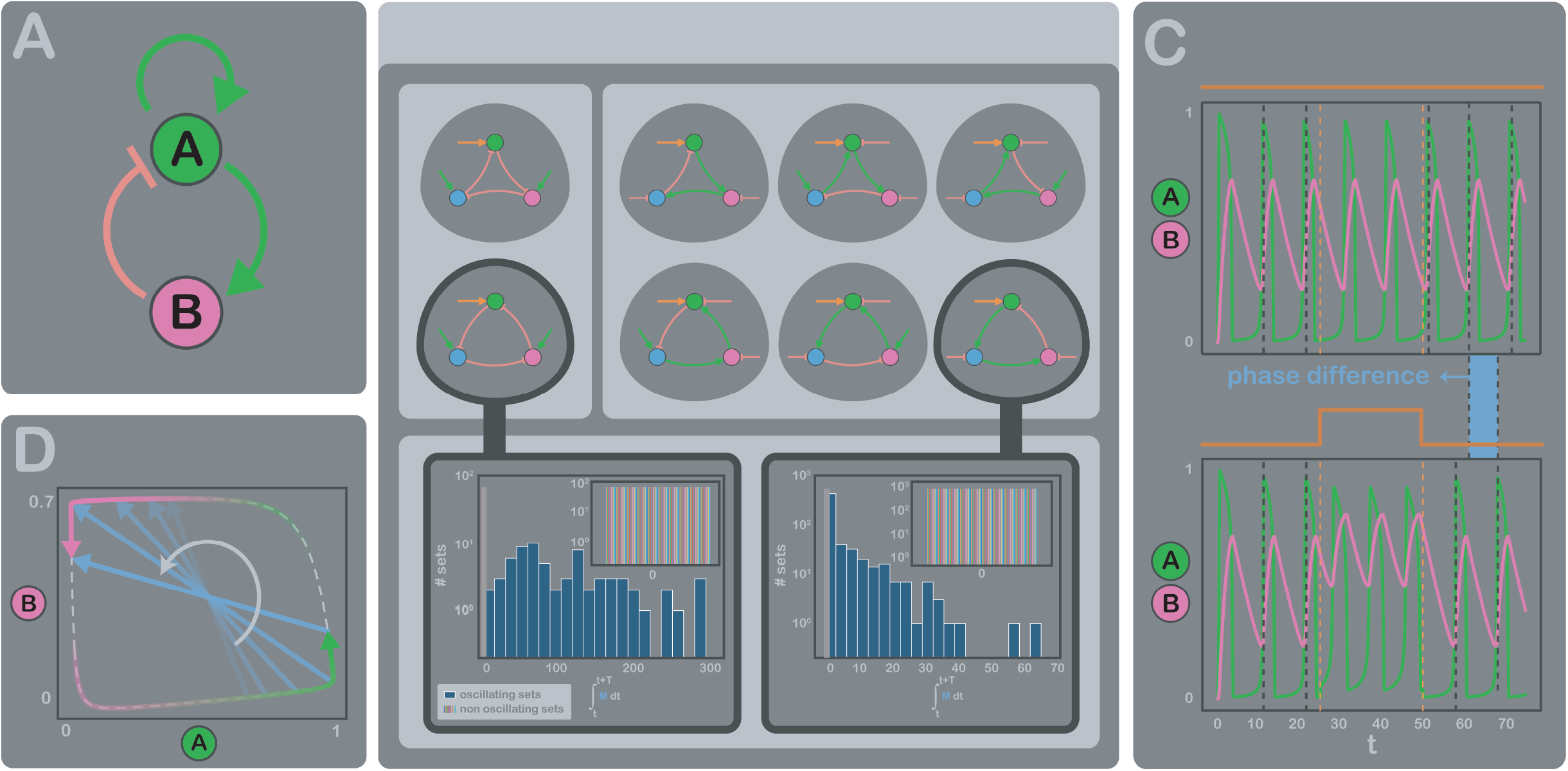
memory in oscillating circuits. A. The simplest circuit which can oscillate. It has a negative and positive feedbacks coupling. Positive feedback provides an hysteresis that introduces the necessary delay to oscillate. The circuit has memory because the stimulus can irreversibly turn on the positive feedback. This memory does not rely on two stable fixed points but is a kind of bistability between a stable fixed point and a limit cycle. B. All 3-node circuits which can oscillate and lack positive feedback. The delay is provided by a third node. The 3-node negative feedbacks can be negative-negative-negative (left panel) or positive-positive-negative (right panel) regulations. Upper circuits are analogous to bottom circuits in clock – anticlockwise symmetry. In addition, positive-positive-negative circuits has symmetry of rotating nodes. Summarizing, each group consists of only one mechanism and so we take both circuits highlighted in black. Bottom panel shows memory distribution of all oscillating sets from 10000 sets of integrated parameters for each circuit. Insets show distribution of 100 groups of the same number of non-oscillating sets, which were all equal to zero. The results show that oscillations can store memory of a stimulus, even in the absence of a positive feedback. C. Temporal curves for the circuit showed in A with and without stimulus. The transient stimulus, once passed, does not change the shape, amplitude, or period of the oscillations. However, it introduces a phase difference between the dynamics with and without it, proving that a system could store information of a transient stimulus in this phase difference. D. Phase diagram of the dynamics with and without stimulus showed in C. We can see the phase difference introduced by the stimulus in the two trajectories (magenta and green), which oscillate around the same limit cycle but maintaining opposite placements. In this example, 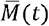 is the rotating vector pointing from one trajectory to another, meaning that memory is stored alternatively in one node and another.

To answer this question, we search for oscillating circuits lacking positive feedbacks. It is known that oscillations need a negative feedback and some delay between the activation of one node and its inhibition (34)(35). This delay can be provided by a third node in the negative feedback, so we take every of our 3-nodes circuits made just of a 3-nodes negative feedback (Figure 6B, upper panels). These circuits can be grouped in two: the repressilators (left) and the positive-positive-negative regulators (right), both widely studied (36)(37)(33). The two repressilators have symmetry of changing clockwise regulations to anti-clockwise. The positive-positive-negative regulators have this same symmetry plus the rotation in the three regulation arrows. We take one circuit from each symmetric group and integrate it over 10000 sets of parameters, chosen randomly again with Latin Hypercube Sampling (20). Each set from each circuit was integrated twice, with and without the stimulus, to compute 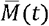 and its integral over a time period.

After integration of both circuits, we automatically labeled parameters sets as oscillating and not oscillating. Figure 6B bottom panel shows distributions of the integral of 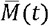 for the oscillating group and for 100 groups of non-oscillating sets. Each set of these groups was randomly chosen from all the non-oscillating, and each group has the same number of sets that the oscillating group. For both circuits, only the oscillating sets have memory, independently of how much. None of non-oscillating sets of all groups has any memory larger than zero (Figure 6C insets). This allows the conclusion that the possibility of oscillating provides memory to a system. Even if it lacks a positive feedback.

This result drives us to the following question. We have proved that an oscillating circuit can remember a given stimulus. Where is this information stored? What can we measure in these circuits to get the information? To answer this, we show the dynamics for an oscillating set (Figure 6C). The comparison between non-stimulated (up) and stimulated dynamics (down) show that the stimulus leaves the oscillations with the same period and amplitude, as there is no bistability and the limit cycle remains intact. However, it does change the phase of oscillations, introducing a phase difference which stores information about the stimulus duration. Figure 6D, shows the phase diagram of these same dynamics, where 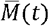 pointing from non-stimulated to stimulated trajectory. In oscillating systems, 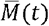 would just remain rotating into the limit cycle. Its module would depend only in the phase difference and shape of the cycle. The phase difference has information of the duration of the stimulus.

In summary, these results suggest that an oscillating system can store memory of a past stimulus in its phase. This could be a hint to understand molecular mechanisms involving cycles. In developmental biology, it has been suggested that stem cells respond to differentiation stimuli only during G1 phase (38)(39)(40). Another coupling founded is between the differentiation of the adipose tissue and the circadian cycle (41)(42)(43). Both are examples of oscillating systems that respond differently to the same stimuli according to its phase. In these cases, a previous priming could change cycle phases and the entire cellular fates.

### mESC remember transient differentiation stimulus

To finish this work, we apply our proposed quantifier to study if mouse Embryonic Stem Cells can remember a differentiation stimulus. We culture these cells in *naïve* state with defined culture medium 2i (29), which maintains the naïve or *ground state* pluripotent identity. The removal of 2i promotes differentiation to a *primed* state (44) where the lineage commitment happens. Figure 7A shows time course experiments we carry on giving a 24h or 48h transient differentiation pulse to naïve cells and test if they go on the differentiation path or they return to naïve state. We test cellular identities by RT-qPCRs for five marker genes: Nanog and Klf4, which express in naïve state, Oct6 and Fgf5, expressed in primed state and Brachyury, a mesendoderm lineage marker. Figure 7B shows fold change of these gene expressions relative to initial naïve state. Nanog and Klf4 both downregulate, as expected, during differentiation pulse, and then partially revert the tendency after pulse end. For Oct6 and Fgf5, we obtained analogous results, but genes first upregulate with the pulse and then downregulate, as expected because they are primed state markers. Our quantifier shows, for these four genes, that cells partially remember the pulse, as they do not completely recover its initial expression values. However, the amount of memory is more 24h post stimulus than 48h after it for all cases, and this might be evidence of some memory loss. For Brachyury, the dynamic is qualitatively different. It upregulates but only once the transient stimulus has ended. This implies that cells are committed to differentiation even when the stimulus has passed, and so they remember it. Our quantifier shows more memory after 48h than 24h, consistently with this commitment.

**Figure 7:**
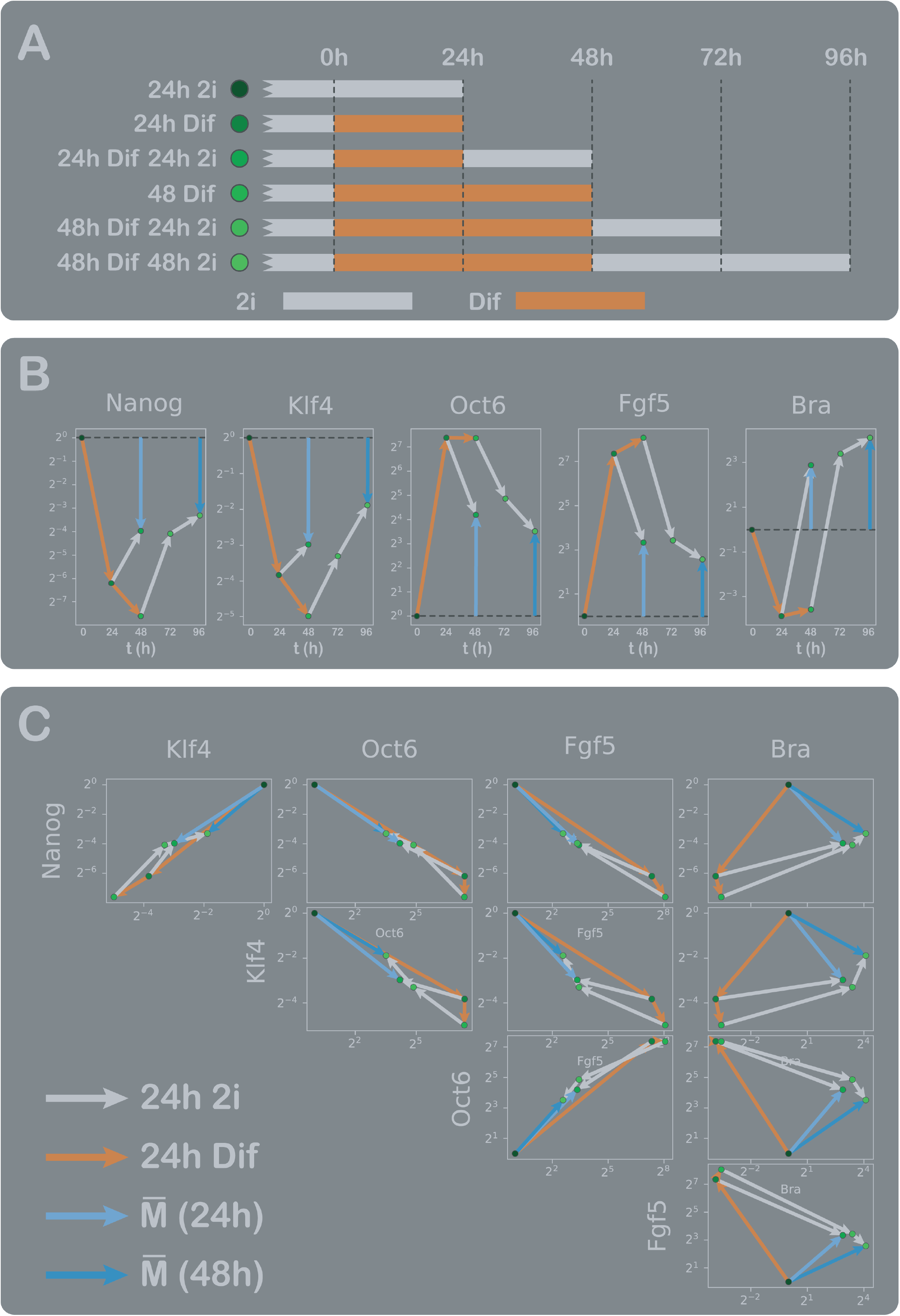
mESC remember differentiation stimulus. A. Time-courses experiments designed to test if mESC remember a transient differentiation stimulus. Light gray indicates 2i medium culture, which maintains naïve pluripotent state. Orange indicates medium culture without 2i, which induces cells to differentiate to primed state and start lineages commitments. We give cells transient differentiation pulses of 24h and 48h and returning times of 24h and 48h. The rest are control conditions to compare. Each condition is labeled in black-green color scale. B. RT-qPCR results for time-course experiments showed in A. Fold change relative to first black condition with only 2i medium. Black to green circles indicate the corresponding time-course and light gray and orange arrows indicate 24hr of culture with medium with or without 2i respectively. Light and dark blue arrows indicate 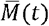 component, corresponding to each gene, 24h or 48h after stimulus respectively. We measure the mRNA expression of two genes associated with the naïve state (Nanog and Klf4), two genes associated with primed state (Oct6 and Fgf5) and one lineage commitment associated gene (Brachyury). C. RT-qPCR results from B, for each pair of genes. Each combination of Nanog, Klf4, Oct6 and Fgf5 indicates that the stimulus takes the system to another state but then, after its removal, it appears to return over the same path. However, Brachyury expression is qualitatively different and shows that cells commit to a different path instead of turning back, which demonstrates they remember the stimulus.

Figure 7C shows the same data as in Figure 7B, but for every pair of genes. We can see that for every combination of Nanog, Klf4, Oct6 and Fgf5, the expression dynamic remains approximately into the same linear path, and goes forward and back. This represents the partial loss of memory, as the expressions tends to go back to the initial point. In contrast, Brachyury expression pulls the dynamic out of a straight line. This reflects the advantage of the vectorial nature of our quantifier. It could happen that memory appears to be lost by just looking at some components of 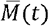 (or some nodes of the system). However, for the whole picture, we need to measure the relevant components, the nodes that are physically responsible to maintain memory. Notice that another advantage is that is not necessary to measure every node of a system. In this example, the measurement of Brachyury alone would be sufficient to determine the system’s memory of the pulse.

## Methods

### Construction of all 1, 2 & 3-nodes circuits combinations

For this study, we analyze every 1, 2 & 3-nodes circuit using the same strategy as previous works (21)(22)(23)(24)(25)(26)(27)(28). We limited to enzymatic regulations using Michaelis-Menten equations. Each node has two reversible states, active (*A*) and inactive (*1-A*), and a fixed total concentration. The interconversion between states is regulated by other node or nodes. Each node can activate, not regulate, or inhibit another, this means, 3 possibilities for each directed link between nodes. 1-node circuits have just the autoregulation link. 2-node have 4, both autoregulation links and one extra link for each node regulating the other. 3-node have 9, 3 links for each node: one for the autoregulation and two for the other nodes regulations. These accounts 3^1^ = 3, 3^4^ = 81 and 3^9^ = 19 683 combinations (3 possibilities for each directed link), but there are some combinations where the output node is disconnected from the input. Discarding these circuits result in 3, 54 and 16038 combinations for 1, 2 & 3-nodes, respectively.

There are some combinations where a node has no incoming activation or inhibition. In these cases, we will assume there is still reversibility by adding a basal activation or inhibition, respectively. For example, the equations for 3-nodes circuits are expressed in (2). The sums are over the specific combination of nodes {*A, B, C*} of each circuit, *x* for the activation links and *Y* for the inhibition ones. There is just one directed link from a node to another: if *A* activates *B* then *A* cannot inhibit *B*. Notice this allows that *B* regulates *A*. Node *A* receives the input so has an additional activation term, and never receives the basal enzyme activation. In nodes of circuits with no incoming links, {*A, B, C*} = Ø, the basal regulation terms 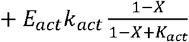 and 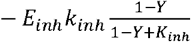 are added for activation and inhibition enzymes, respectively.

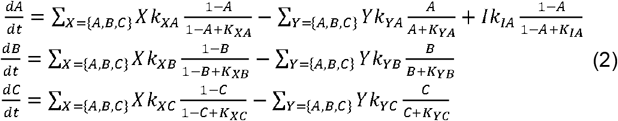

Each term has 2 parameters (*k* and *K*), so each circuit has *2#*_*terms*_ parameters. The minimum number of terms *(#*_*terms*_) is just one activation and inhibition for each node: *2#*_*nodes*_. The input term is counted as the activation of node *A*. The maximum is each node activated or inhibited by itself and all the other ones plus one basal enzyme for each node plus the input: 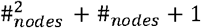. In summary, number of parameters ranges from 4-6 for 1-node, 8-14 for 2-nodes and 12 - 26 for 3-nodes circuits. We take an exhaustive parameter scanning of 10000 random sets for each circuit topology, chosen by Latin Hypercube Sampling (20). Each parameter has a log-uniform distribution, every *k* ranges from 10^−1^ to 10^1^ and every *K* from 10^−2^ to 10^2^. In total, we have integrated (3+ 54+ 16038) x 10 000 = 160 950 000 different circuits for this study.

### Numerical integration of circuits and memory computation

We compute memory by integrating each circuit for the 3 stages (low, high, low) of the square pulse input showed in Figure 1 C and 2. For every integration, low value of input was 0.1 and high 5. The initial condition of first stage was zero for all nodes, and the next stages take the previous one final values of each node as the initial conditions, to account for continuity. We use and adaptative step Runge Kutta method in Python to optimize integration of circuits with wide variety of timescales. On each stage, integration continues until steady state was reached. We assume this when every node moves less than 10^−5^ from one step to another. This assumption fails when circuits have nodes with very long timescales (≈ 1% cases). In these cases, integration can finish before steady state is reached, but increasing this threshold makes every integration longer. The 10^−5^ threshold was chosen to minimize these incorrect cases but keeping the integration time short enough to accomplish the 160 950 000 circuits solving in a reasonable time.

We compute long-term memory of each set of parameters of each circuit topology as indicated in equation (5). 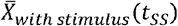 is the vector of 1, 2 or 3 nodes values at the final time of the third stage of integration. 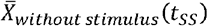 is the value at the final time of the first stage of integration. Notice this is equivalent to the steady state of the circuit with no square pulse stimulus but saves us from integrating each circuit again.

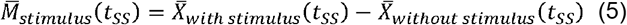

We saved these vectors for all circuits, and they compose our whole dataset illustrated in Figure 3B table.

### Choice of arbitrary thresholds to determine if a circuit topology has long-term memory

Figure 2 illustrates the algorithm designed to determine if a given circuit topology has long-term memory. This determination depends finally on two different thresholds, showed in Figure 2 histograms of right panels. The aim of these thresholds is to discard slower circuits with 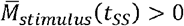 measured. This memory can be due to an interrupted integration before the steady state is reached, instead of a real memory due to bistability or other mechanisms.

To characterize this artifact in memory computation, we take two simple circuits topologies showed in Figure 2 a and B, left panels. Panel A topology has no memory and panel B has, because it can have bistability for some sets of parameters. We measure 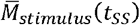 distributions for the 10 000 sets of parameters of each and then we labeled each set according to the integration reached steady state or was interrupted before. These labels where defined by just taking the final state of circuits and adding time steps to the integration. This operation takes a long time for all *160 950 000* circuits but we can afford it for just two topologies. For each topology, we have one distribution divided in two groups: steady state circuits and interrupted ones. Results are showed in histograms of right panels in Figure 2 A and B. As expected, there is no complete integrated circuit in panel A with memory, as its topology does not have bistability. Every circuit with 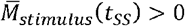 is just an artifact of interrupted integration. Comparing with panel B distribution, we can see that there are completely integrated circuits with 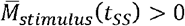, which corresponds to real memory due to the bistability.

In general, we have just the sum of both labeled distributions, where there are circuits with 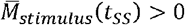 but no real memory. Each circuit with no memory, even the slower ones, must accomplish that 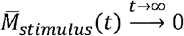. So, these artificial cases have low probability of 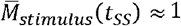. *T*_*M*_ was chosen from panel A circuit distribution, the criteria was to take the minimum value greater than every 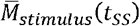 due to this artifact on this topology. This value resulted in *T*_*M*_ = 0.8.

We choose *T*_*M*_ value from just one topology. To be stricter in the definition of all topologies with memory, we add an additional threshold *T*_*sets*_. This is for the number of sets of parameters. A given topology must have more than *T*_*sets*_ circuits where 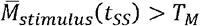 to determine that it has long-term memory. This threshold was fixed in *T*_*sets*_ = 3, and was also inspired in Figure 2 histograms. In general, if a given topology has bistability, there will be plenty of sets with 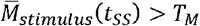, as showed in Figure 2 B. In contrast, by analyzing more topologies with no memory at all we see that there are never more than *2* circuits where 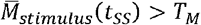.

These two thresholds allow us to filter circuits with real memory and strictly define that a topology has long-term memory. Both are showed in the two histograms of Figure 2 A and B right panels. Every topology with long-term memory has values in the marked rectangular region.

### Neighboring circuits comparative algorithm

The aim of this algorithm is to quantify how good or bad is a motif to provide a given circuit with some quantifiable property. Here we are interested in memory, measured as indicated in equation (5). The algorithm takes a given motif as the input and returns a number indicating how much more or less memory this motif provides to all circuits having it. We call this number *CNM* for *comparative neighboring memory*, and it is calculated as follows. First, we take the desired motif. Second, we go to the corresponding meta-network and get the pairs of neighboring circuits. We did this step computationally, filtering all circuits with the motif and then, on each circuit, removing one regulation of the motif at a time to get it neighbor pair. This way, each pair contains one circuit with the motif and another similar one but with the motif broken. Figure 5B illustrates a schematic example of three pairs of circuits where the motif is the positive autoregulation of the magenta node. Third, sum the memory of all circuits for the 10000 sets of parameters. We decided to take just one component of 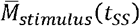, thinking it as the memory stored in just the output (magenta) node. Finally, for each neighbor pair compute the difference between the sum from the circuits with the motif and the sets from circuits without the motif and get the average for all pairs.Equation (6) summarize last two steps.

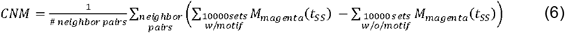

Notice that *CNM* can be positive or negative. A motif having positive *CNM* means that, on average, it provides memory to a circuit. Negative *CNM* means it removes memory and *CNM* near zero means the motif is not related to memory.

To understand how much different from zero must be any *CNM* to relate a motif to memory, we design a control measure. It consists of taking the sum over neighbor pairs and interchange both terms with 50% of probability on each one. We repeat this process 10000 times for each motif to get a distribution of control *CNMs*. This distribution has to be centered in zero, but it allows us to compare its deviation with the original *CNM*. Figure 5 C and D show the 50% and 95% confidence intervals from these distributions in orange bands. Bars over this bands means the motif really provides memory to a circuit and, down this bands, that really erases it.

## Discussion

The presented work focuses on developing a general mathematical framework for defining and quantifying memory in complex dynamical networks, with a particular emphasis on gene regulatory networks. The key contributions and findings of the study are discussed below.

### Definition of Memory

The introduction of a memory quantifier provides a unique, comparable, and comprehensive measure for capturing the memory of a general dynamic system in response to a transient stimulus. This quantifier is derived by comparing the system’s state with and without the stimulus over time.

### Validation Through Toy Models

The proposed memory quantifier was validated using toy models, specifically a two-node system with positive feedback loops. The examples demonstrate how bistability in positive feedback loops results in memory, emphasizing the role of stable fixed points and hysteresis.

### Algorithm for Long-Term Memory and minimal Motifs results

We introduced an algorithm to determine if any dynamical system exhibits long-term memory. It involves the computation for various parameter sets and define thresholds to classify circuits based on their memory.

The study identifies minimal motifs responsible for providing long-term memory in gene regulatory circuits. The findings indicate that positive feedback loops play a crucial role in the mechanisms that enable memory. Minimal motifs for one, two, and three-node circuits are illustrated and discussed.

These findings are consistent with some previous evidence found in the literature. The algorithm serves as a ground truth for researching the memory of even more complex systems. In particular, the discovered symmetry of interchange between activating and inactivating nodes could be generalized to other computational functions, such as adaptation or fold-change response. The main idea is that, for each circuit exhibiting the function, there exists another symmetrical circuit that also possesses it.

### Comparison of Circuits with Positive and Negative Feedback

The research introduces a comparative algorithm to analyze whether circuits with positive feedback have more memory than those without. The results show that positive feedback enhances memory, while certain negative feedbacks can diminish it due to a shortened dynamic range.

Beyond these findings, the neighbor circuits comparative algorithm can be applied in the context of any motifs studied and also with any computational function or dynamical property of interest. It responds the more general question if a given motif enhances or diminish some feature.

### Oscillations and Memory

The study explores the relationship between oscillations and memory. We demonstrated that oscillating circuits, even without positive feedback, can exhibit memory. This consist of an entirely different mechanism of storing memory, as it lacks any positive feedback. The main idea is that the stimulus can introduce a delay in oscillations. Then, the phase difference is the key parameter storing information about the duration of a stimulus.

Looking at steady state dynamics, oscillations are just the next step in complexity from fixed points. The fact that they can store memory is a very interesting example of how many computational properties we can expect from even complex systems with more nodes or non-linear effects.

### Experimental Validation in mESCs

We applied the proposed memory quantifier to experimental data involving mouse Embryonic Stem Cells (mESCs) subjected to transient differentiation stimuli. The results show that mESCs retain memory of the stimulus, with different genes exhibiting varying degrees of memory.

The study emphasizes the vectorial nature of the proposed memory quantifier, showcasing its advantage in capturing the system’s memory holistically. This is illustrated both through the analysis of gene expression dynamics in mESCs and the toy models of two and three nodes.

### Implications for Biological Systems

The findings have potential implications for understanding molecular mechanisms, especially in developmental biology. The study suggests that memory in biological systems may not solely rely on bistability but can also be associated with oscillatory behavior.

In conclusion, the work presents a comprehensive framework for defining, quantifying, and studying memory in complex dynamical networks, with applications in gene regulatory circuits. The combination of mathematical modeling, computational analysis, and experimental validation contributes to a deeper understanding of memory mechanisms in biological systems.

## Captions

**Figure S1:**
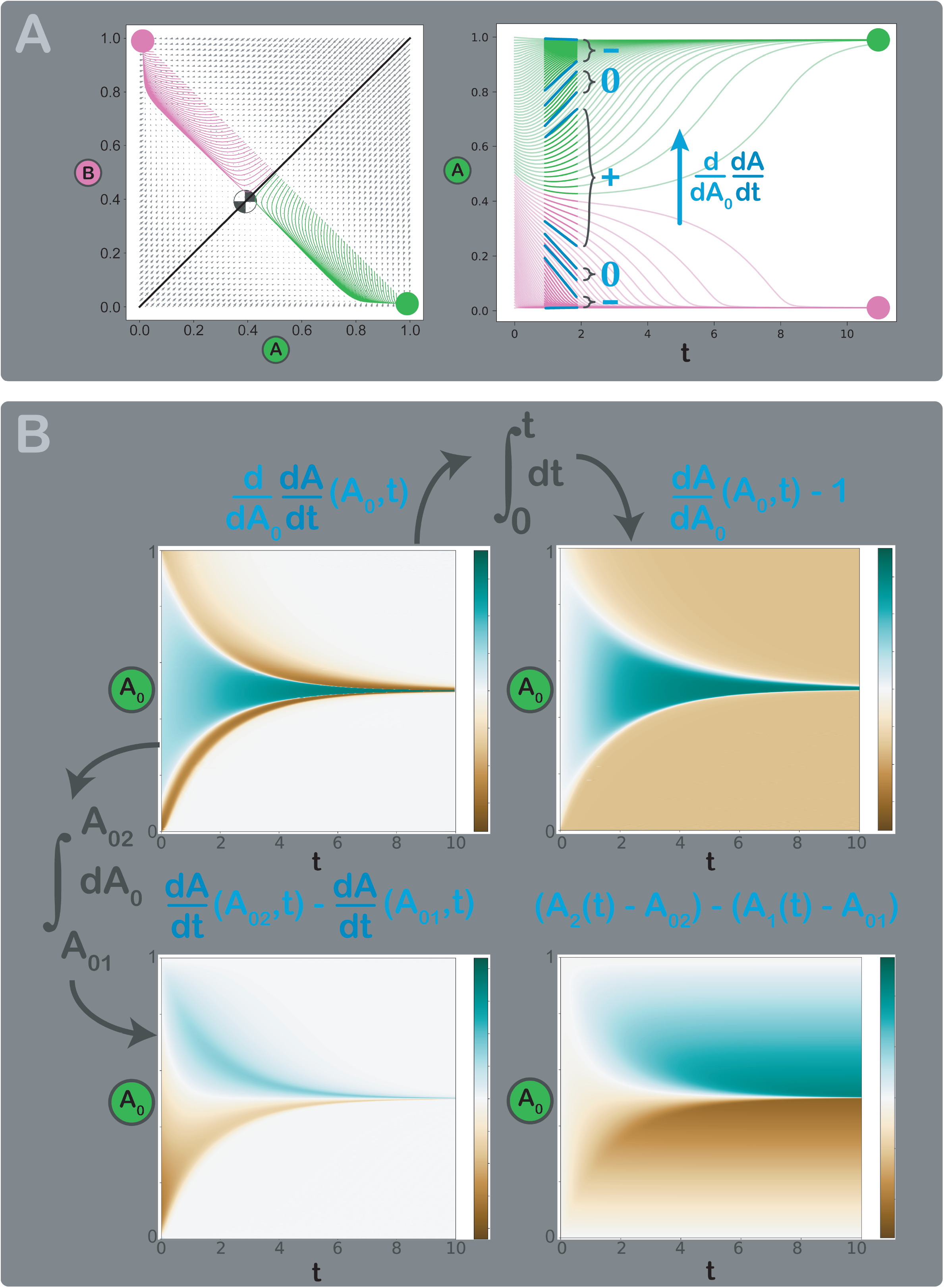
Example of relation between memory quantifier and information retained. A. Phase space trajectories for the model in Figure 1A, with a sweep of initial conditions from high A and low B to low A and high B (left panel). Dynamics of node A for the same trajectories. We can observe that, in the vicinity of the saddle point, trajectories diverge, while in the vicinity of stable points, they converge. It is possible to quantify this convergence or separation as the change in the slope of trajectories as we move through initial conditions (S1). B. Transition between convergence or separation and the memory quantifier. In the top left panel, the change in slopes (color-coded) over time and initial condition is depicted. Notice the positive or greener values in trajectories when diverging, a zone that extends in time only in the narrow region near the saddle point. Both orange and negative values surrounding the green region represent the time of convergence of those trajectories to both stable fixed points. Once the convergence has passed, the white zone represents the lack of convergence or divergence as the trajectories remain still. Moving to the right, integrating with respect to time, we obtain the net convergence between initial time t = 0 and final time t = 10 ((S2), upper right panel). Notice that almost all trajectories finish with negative orange values, which means they have a net convergence within their own vicinity. The exception is just the vicinity of *A*_*0*_ = 0.5 where curves diverge to both different stable fixed points. Moving downward, integrating with respect to the initial value *A*_*0*_, we obtain the net convergence between two different trajectories ((S3), bottom left panel). We took *A*_*0*_ =0 as the reference trajectory, and the color-graph indicates how much a trajectory is converging or diverging from this one. During some memory measurement, this reference trajectory would be our system without stimulus. Green zones represent curves when diverging from the reference trajectory, and orange ones represent curves when converging. The bottom right panel shows the memory quantifier, a result of the double integration ((S4) or (S5)). From the left panel, we just take the net convergence to the reference trajectory in time. The graph divides into two different zones. The orange zone corresponds to trajectories converging to the same fixed point as the reference, so they have a net loss of memory. The green zone corresponds to trajectories converging to the other fixed point, so they are distinguishable from the reference and have retained information.

