## Supplementary material for "Defining Network Topologies that Can Achieve Molecular Memory": Suplementary Text

### Supplementary Text

In the introduction of this work, we presented different definitions of memory and discussed their relations and equivalences. In this section, we will mathematically prove that the amount of information retained from the past is exactly what our proposed quantifier measures. It is mathematically equivalent to the difference between a system’s state with and without having received a transient stimulus.

Imagine that we want to measure if a system has received a transient stimulus. We need the system’s state, during the measurement, to be sufficiently different from its normal state without the stimulus. "Sufficiently different" is according to our measurement resolution. This means that if some measured variable, in the stimulated state, is too close to its normal value without the stimulation, we would not be able to distinguish between both states. This was the initial motivation to quantify how much two different states converge or diverge from each other in time.

The time derivative of the expression in eq. (S1) quantifies the slope, at a given time, of some dynamic variable $A$. The second derivative with respect to $A_{0}$, the initial condition, gives the rate of change of the slope of each curve moving through the initial values.

$\frac{d}{dA_{0}}\frac{dA}{dt}\left( A_{0},t \right)$	(S1)

Figure S1**A** illustrates an example of this quantifier application to our double negative regulation toy model. Looking at some given time and some given $A_{0}$, it shows that our quantifier would be positive, zero, or negative depending on if the curves are diverging, being parallel, or converging from each other in the surroundings of $A_{0}$. Relating this with our experimental resolution, we are quantifying how much information we are losing from the initial condition.

The expression in eq. (S1) provides a local quantifier of how much information we are gaining or losing at a given time point and from a given initial condition. It is not really useful if we want to know how much is lost during an entire experiment. However, we can integrate the expression in time to get the total convergence or divergence of curves from a given initial time, like the time when the stimulus starts, to a final time when the measurement is done.

$\int_{t_{0}}^{t} \frac{d}{dA_{0}}\frac{dA}{dt'}\left( A_{0},t' \right)dt^{'}=\frac{dA}{dA_{0}}(A_{0},t)-\frac{dA}{dA_{0}}(A_{0},t_{0}) =\frac{dA}{dA_{0}}(A_{0},t)-1$	(S2)

Eq. (S2) quantifies how much net convergence or divergence is from an initial time $t_{0}$ to a final time $t$ in the surroundings of $A_{0}$.

Analogously, we can integrate eq. (S1) with respect to $A_{0}$ and obtain the net convergence or divergence at a given time point between two separate initial conditions $A_{01}$ and $A_{02}$ (eq. (S3)):

$\int_{A_{01}}^{A_{02}} \frac{d}{dA_{0}}\frac{dA}{dt}\left( A_{0},t \right)dA_{0}=\frac{dA}{dt}(A_{02},t)-\frac{dA}{dt}(A_{01},t)$	(S3)

Finally, as $A_{0}$ and $t$ are independent variables, we can integrate both eq. (S2) with respect to $t$ or eq. (S3) with respect to $A_{0}$. Eq. (S4) shows the integration of (S2), where we consider that $A\left( A_{02},t_{0} \right)-A\left( A_{01},t_{0} \right)=A_{02}-A_{01}=0$. The last equality is because both initial conditions are before the stimulus is given, where curves depart from the same place.

$$\int_{A_{01}}^{A_{02}} \left[ \frac{dA}{dA_{0}}\left( A_{0},t \right)-1 \right]dA_{0}=A\left( A_{02},t \right)-A\left( A_{02},t_{0} \right)-\left[ A\left( A_{01},t \right)-A\left( A_{01},t_{0} \right) \right]$$

$\int_{A_{01}}^{A_{02}} \left[ \frac{dA}{dA_{0}}\left( A_{0},t \right)-1 \right]dA_{0}=A\left( A_{02},t \right)-A\left( A_{01},t \right)$	(S4)

Eq. (S5) shows the integration of (S3):

$\int_{t_{0}}^{t} \left[ \frac{dA}{dt^{'}}\left( A_{02},t^{'} \right)-\frac{dA}{dt^{'}}\left( A_{01},t^{'} \right) \right]dt^{'}=A\left( A_{02},t \right)-A\left( A_{01},t \right)$	(S5)

This concludes our proposed memory quantifier, which provides the total convergence or divergence of two curves departing from $A_{01}$ and $A_{02}$, from an initial time $t_{0}$ to a final time where the measurement takes place. The difference between (S5) and (1) is that $A$ represents just one measured variable of a system; it would be one coordinate of the vector $\bar{M}$. In addition, we can think of both curves from $A_{01}$ and $A_{02}$ as the system with and without the transient stimulus.

Figure S1**B** illustrates all mathematical quantities (S1), (S2), (S3), and our proposed memory quantifier of (S4) and (S5). It shows the application of these measurements to our double negative regulation toy model. As an example, we took the same scanning of $A_{0}$ curves from Figure S1**A**, from (1,0) to (0,1) as initial conditions. Notice the deep relation between the convergence or divergence of curves (Figure S1**A**) and the final memory measured (Figure S1**B**).

Summarizing, we have mathematically related the idea of a system retaining information from the past to our proposed memory quantifier. Taking the difference between a system state with and without a stimulus is equivalent to measuring how much those two dynamics converge to each other during an experimental time window.
